# Antimicrobial Susceptibility and Genomic Determinants of Resistance and Virulence in *Mycoplasma cynos* and *Mycoplasma felis*

**DOI:** 10.1101/2025.06.02.657554

**Authors:** Isaac Framst, Michael L. Beeton, Shelley W. Peterson, Irene Martin, Jeff Caswell, Grazieli Maboni

## Abstract

*Mycoplasma cynos* and *Mycoplasma felis* are important respiratory pathogens in dogs and cats. Due to the challenges of culturing these fastidious bacteria, little is known about their antimicrobial susceptibility or mechanisms of pathogenicity. Treatment is typically empirical, as *in vitro* antimicrobial activity has not been evaluated, and therapeutic efficacy remains unclear. This study aimed to assess *in vitro* susceptibility and identify genetic markers of antimicrobial resistance (AMR) and virulence in *M. cynos* and *M. felis* clinical isolates. Minimum inhibitory concentrations (MICs) for doxycycline, tetracycline, minocycline, enrofloxacin, marbofloxacin, and azithromycin were determined using a broth microdilution assay developed for this study. Hybrid genomes were generated using Oxford Nanopore and Illumina sequencing. AMR-associated single-nucleotide polymorphisms (SNPs) in the *gyrA* gene correlated with high MICs to enrofloxacin and marbofloxacin in both species. Mutations in 23S rRNA were associated with reduced susceptibility to azithromycin. In *M. felis*, novel variants in *gyrA* and the 50S ribosomal protein L4 were linked to decreased susceptibility to fluoroquinolones and azithromycin, respectively. The data also suggest potential intrinsic resistance to azithromycin in *M. felis*. All isolates required low MICs of tetracyclines, and resistance mutations were not identified in the 16S rRNA gene, supporting tetracyclines as effective first-line treatment options. Virulence genes, particularly those associated with adhesion and immune evasion, were detected in both *M. cynos* and *M. felis*. This study presents the first comprehensive genomic and phenotypic analysis of AMR and virulence in *M. cynos* and *M. felis*, providing new insights into their pathogenicity and informing evidence-based therapeutic strategies.

## Introduction

*Mycoplasma cynos* and *Mycoplasma felis* are increasingly recognized as significant respiratory pathogens in dogs and cats, respectively. They are highly fastidious and difficult to culture, which likely contributes to the limited research on their pathogenicity, antimicrobial susceptibility, and the scarcity of diagnostic tools.

*M. cynos* and *M. felis* are commonly associated with canine infectious respiratory disease complex (CIRDC) and feline respiratory disease complex (FRDC), respectively (Day et al., 2020; Le Boedec, 2017; Maboni et al., 2019; Nguyen et al., 2019; Priestnall et al., 2014). CIRDC, also known as "kennel cough," can be highly contagious and presents with various clinical signs ranging from mild cough and sneezing to tracheobronchitis and fatal pneumonia (Day et al., 2020; Maboni et al., 2019; Voorhees et al., 2017). *Mycoplasma cynos* is a proven causative agent of CIRDC (Rosendal & Vinther, 1977), and epidemiological studies in North America and Europe revealed that *M. cynos* was among the most common pathogens associated with CIRDC (Chalker et al., 2004; Day et al., 2020; Jambhekar et al., 2019; Maboni et al., 2019; Yondo et al., 2023). Feline Respiratory Disease Complex (FRDC) is a contagious respiratory infection in cats with upper and lower respiratory signs, conjunctivitis, and stomatitis (Lee-Fowler, 2014; Nguyen et al., 2019). As for CIRDC, recent studies conducted in Australia, USA, and Spain reported that *M. felis* was among the most common pathogens associated with FRDC (Fernandez et al., 2017; Maboni et al., 2024; Nguyen et al., 2019).

The current treatment for *Mycoplasma*-associated respiratory infections in dogs and cats relies on the empirical use of antimicrobials including tetracyclines (e.g., doxycycline, tetracycline, and minocycline), fluoroquinolones (e.g., marbofloxacin, enrofloxacin, and pradofloxacin), and macrolides (e.g. azithromycin) (Lappin et al., 2017). There are no published *in vitro* susceptibility studies for *M. cynos* and *M. felis*, hence, the inhibitory or bactericidal effect of antimicrobials is unknown, and the effectiveness of therapy is not well documented (Abd El-Hamid et al., 2019; Lappin et al., 2017; Lysnyansky & Ayling, 2016). Additionally, due to the lack of a cell wall and the absence of a metabolic pathway to synthesize folate, mycoplasmas are intrinsically resistant to beta-lactams and sulfonamides (Maunsell et al., 2011). Furthermore, fluoroquinolone and macrolide resistance have been commonly reported in other respiratory *Mycoplasma* species such as *M. pneumoniae* (Dong et al., 2022) and *M. amphoriforme* in humans (Day et al., 2022), *M. bovis* in cattle (Bokma et al., 2021), *M. ovipneumoniae* in bighorn sheep (Spaan et al., 2021), *M. hyopneumoniae* in swine (Liu et al., 2019), and *M. gallisepticum* in poultry (Feberwee et al., 2022).

We aimed to evaluate the *in vitro* activity of antimicrobials against clinical isolates of *M. cynos* and *M. felis.* To achieve this, we developed a broth microdilution method to determine the minimum inhibitory concentration (MIC) of selected antimicrobial agents. A second aim was to identify genetic markers linked to antimicrobial resistance and virulence in *M. cynos* and *M. felis* hybrid whole genome sequences.

## Methods

### Study population

This retrospective study utilized samples submitted by licensed veterinarians from client-owned dogs and cats to the Mycoplasmology service at the Animal Health Laboratory (AHL, Ontario Veterinary College, ON, Canada) for the diagnosis of *Mycoplasma*-related respiratory infections between 2015 and 2022. Clinical samples were accompanied by a submission form with information about the clinical history, vaccination records, and clinical signs. These forms were retrieved from the AHL electronic database, and for each isolate, information about the month, year, animal species, sex, age, clinical history, and clinical specimen was recorded (Table 1).

**Table 1.** Sample and whole genome assembly information of *Mycoplasma cynos* and *Mycoplasma felis* clinical isolates investigated in this study.

| Isolate ID<sup>a</sup> | Species | Isolate<sup>b</sup> | Host | Isolation Source | Year | SRA accession |  | Genome Accession |
| --- | --- | --- | --- | --- | --- | --- | --- | --- |
|  |  |  |  |  |  | ONT | Illumina |  |
| C01 | *M. cynos* | 30510 | *Canis lupus familiaris* | Bronchoalveolar Lavage | 2018 | SRR26841336 | SRR27086663 | CP141046 |
| C02 | *M. cynos* | 78726 | *Canis lupus familiaris* | Transtracheal wash | 2016 | SRR26841316 | SRR27086662 | CP141045 |
| C03 | *M. cynos* | 98999 | *Canis lupus familiaris* | Bronchoalveolar Lavage | 2017 | SRR26841315 | SRR27086651 | CP141044 |
| C04 | *M. cynos* | 97732 | *Canis lupus familiaris* | Bronchoalveolar Lavage | 2015 | SRR26841314 | SRR27086645 | CP141043 |
| C06 | *M. cynos* | 80950 | *Canis lupus familiaris* | Bronchoalveolar Lavage | 2015 | SRR26841313 | SRR27086644 | CP141042 |
| C07 | *M. cynos* | 87299 | *Canis lupus familiaris* | Nasal swab | 2018 | SRR27145255 | SRR27145254 | CP140753 |
| C09 | *M. cynos* | 94662 | *Canis lupus familiaris* | Bronchoalveolar Lavage | 2017 | SRR26841312 | SRR27086643 | CP141041 |
| C10 | *M. cynos* | 85201 | *Canis lupus familiaris* | Bronchoalveolar Lavage | 2021 | SRR26841335 | SRR27086642 | CP141040 |
| C11 | *M. cynos* | 71053 | *Canis lupus familiaris* | Bronchoalveolar Lavage | 2021 | SRR26841324 | SRR27086641 | CP141039 |
| C12 | *M. cynos* | 64228 | *Canis lupus familiaris* | Nasal swab | 2020 | SRR26841318 | SRR27086640 | CP141038 |
| C14 | *M. cynos* | 58334 | *Canis lupus familiaris* | Urine | 2021 | SRR26841317 | SRR27086639 | CP141036 |
| F01 | *M. felis* | 85384 | *Felis catus* | Transtracheal wash | 2016 | SRR26841334 | SRR27086661 | CP141035 |
| F02 | *M. felis* | 21976 | *Felis catus* | Nasal swab | 2022 | SRR26841326 | SRR27086660 | CP141034 |
| F03 | *M. felis* | 35610 | *Felis catus* | Nasal swab | 2022 | SRR26841325 | SRR27086659 | CP141033 |
| F04 | *M. felis* | 40612 | *Felis catus* | Bronchoalveolar Lavage | 2016 | SRR26841323 | SRR27086658 | CP141032 |
| F06 | *M. felis* | 57065 | *Felis catus* | Bronchoalveolar Lavage | 2016 | SRR26841322 | SRR27086657 | CP141031 |
| F07 | *M. felis* | 79979 | *Felis catus* | Bronchoalveolar Lavage | 2015 | SRR26841321 | SRR27086656 | CP141030 |
| F08* | *M. felis* | 99751 | *Felis catus* | Nasal swab | 2017 | SRR26841320 | SRR27086655 | CP141029 |
| F09 | *M. felis* | 11534 | *Felis catus* | Transtracheal wash | 2017 | SRR26841319 | SRR27086654 | CP141028 |
| F10 | *M. felis* | 53109 | *Felis catus* | Bronchoalveolar Lavage | 2020 | SRR26841333 | SRR27086653 | CP141047 |
| F11 | *M. felis* | 99014 | *Felis catus* | Nasal swab | 2016 | SRR26841332 | SRR27086652 | CP141027 |
| F13 | *M. felis* | 64570 | *Felis catus* | Nasal swab | 2020 | SRR26841330 | SRR27086649 | CP141025 |
| F14 | *M. felis* | 71855 | *Felis catus* | Bronchoalveolar Lavage | 2021 | SRR26841329 | SRR27086648 | CP141024 |
| F15 | *M. felis* | 27649 | *Felis catus* | N/A | 2021 | SRR26841328 | SRR27086647 | CP141023 |
| F16 | *M. felis* | 04555 | *Felis catus* | Transtracheal wash | 2022 | SRR26841327 | SRR27086646 | CP141036 |

\*Reference genome for *Mycoplasma felis*
<sup>a</sup>In-house sample number. C: *Mycoplasma cynos*; F: *Mycoplasma felis*
<sup>b</sup>Isolate tracking number used by the Animal Health Laboratory, Guelph, Ontario, Canada
N/A: information not available

*Mycoplasma* genus and species identification were part of the AHL diagnostic work-up using agar culture methods and fluorescent antibody testing. Selected isolates were then cryopreserved at -80 °C. *Mycoplasmopsis* is now considered a homotypic synonym of the genus *Mycoplasma*, and this term was accepted by the *International Journal of Systematic and Evolutionary Microbiology* (May & Brown, 2019; Munson et al., 2023).

### Colour-changing unit (CCU) assay for *M. cynos* and *M. felis*

Inoculum preparation of *M. cynos* and *M. felis* was based on CCU following the M43-A CLSI guidelines and the method described by Hannan *et al*., 2000, with modifications. First, two media were evaluated for the optimal growth of *M. cynos* and *M. felis*: a commercial medium, SP4 broth with glucose (Hardy Diagnostics: U86), and a medium optimized in-house, named “super-modified Hayflick’s broth” (**Supplementary file 1**). CCU assays were performed in duplicate in 96-well plates, and the inoculum concentration was determined as described in the M43-A CLSI protocol. CCU assays were repeated twice for each isolate. *M. cynos* CCU assays were incubated at 37 °C with 5% CO_2_ for 72 h, while *M. felis* assays were incubated at 37 °C with 5% CO_2_ for 48 h. Growth was visualized using 5% AlamarBlue (ThermoFisher Scientific).

### Broth microdilution assay for *M. cynos* and *M. felis*

Tetracycline, doxycycline, minocycline, azithromycin, marbofloxacin, and enrofloxacin (Sigma-Aldrich) were prepared according to the manufacturer’s instructions and CLSI M43 guidelines (Clinical and Laboratory Standards Institute (CLSI), 2011). The broth microdilution assay was then performed with an inoculum concentration ranging from 10^3^ to 10^5^ CCU. The inoculum was diluted or concentrated to achieve a final concentration between 10^^3^ and 10^^5^ CCU. To reduce cell clumping, the inoculum of each isolate was passed through 2 mm needles. Broth microdilution assays were performed using 96-well microdilution plates according to the CLSI document M34-A (CLSI, 2011). *M. cynos* assays were incubated for 72 h and *M. felis* for 48 h at 37 °C with 5% CO_2_.

A *M. cynos* reference strain isolated by Rosendal *et al*. in 1973, purchased from the American Type Culture Collection (ATCC 27544 – Rosendal strain H831, 1973, NCBI BioSample SAMEA4412697), was included in this study. *Mycoplasmopsis bovis* strain PG45, with well-characterized minimum inhibitory concentrations (MICs), was tested to compare our results with those of other publications.

The MIC was defined as the “lowest concentration of an antimicrobial that inhibited the visible growth of *M. cynos* or *M. felis* after incubation,” visualized by no colour change in the well (Hannan, 2000). When the highest antimicrobial concentration did not inhibit growth, the MIC was expressed as equal to or greater than (≥) the highest antibiotic concentration tested. Similarly, if growth was inhibited by the lowest antibiotic concentration in the plate, the MIC value was expressed as less than or equal to (≤) that concentration.

### Whole genome sequencing

DNA extraction, library preparation, and Oxford Nanopore Technologies (ONT) long-read sequencing for *M. cynos* and *M. felis* were performed as described previously by our group (Framst et al., 2022). Illumina short-read sequencing was conducted by a third-party facility using the Nextera DNA Flex kit with 15x sequencing depth (Advanced Analysis Center Genomics, University of Guelph, Guelph, Ontario, Canada).

Hybrid *de novo* whole genome assembly of *M. cynos* and *M. felis* was performed as described previously by our group (Framst et al., 2024). The resultant assemblies were verified as *M. cynos* or *M. felis* species by BLAST search of the 16S-23S intergenic spacer region. Assembly quality and completeness were assessed using QUAST v5.2.0 (https://github.com/ablab/quast) (Gurevich et al., 2013). Genomes were evaluated for quality and completeness using BUSCO v5.7.0 (https://busco.ezlab.org) with the mycoplasmatales_odb10 database (Manni et al., 2021).

### Identification of antimicrobial resistance mutations

Gene sequences were extracted using the MasterBlastR function of the WGS Analysis and Detection of Molecular Markers (WADE) program (https://github.com/phac-nml/wade) and verified using Geneious v2023.0. The extracted genes of interest were aligned using MUSCLE and compared against genes from *M. cynos* C142 and *M. felis* Myco-2 reference strains using Aliview v1.28 (https://ormbunkar.se/aliview/). For genes not annotated in the *M. cynos* C142 and the *M. felis* Myco-2 strains, the conserved region of the gene of interest in the *Mycoplasmopsis bovis* PG45 (NC_014760) was used as a template to find and extract the gene of interest in the reference *M. cynos* C142 or *M. felis* Myco-2 genome. Gene sequences were manually aligned, and SNPs were detected using AliView (https://github.com/AliView/AliView) (Larsson, 2014). Coding sequences were translated and reading frame shifts were adjusted to align the amino acid sequences. Single nucleotide polymorphisms (SNPs) previously associated with antimicrobial resistance in *E. coli* were evaluated. Specifically, tetracycline resistance-conferring SNPs A965, A967, G1058, and C1192 in the 16S rRNA gene were mapped to positions A928, A930, G1017, and C1155 in *M. cynos*, or A933, A935, G1022, and C1163 in *M. felis*. Similarly, macrolide resistance-conferring positions G748A, A2058G, A2062G, and C2611T in the 23S rRNA gene were mapped to positions G791, A2058, A2062, and C2610 in *M. cynos*, and G790, A2057, A2061, and C2610 in *M. felis*. Additionally, the GyrA peptide positions Ser83Phe and Glu87Gln/Val were mapped to Tyr154Ala/Ile and Asp158 in *M. cynos*, and Ser137Tyr to Glu141Gln/Gly in *M. felis*. COPLA (https://castillo.dicom.unican.es/copla/) was used to identify potential plasmids (Redondo-Salvo et al., 2020), and potential acquisition of resistance genes was screened using the CARD database.

### Identification of virulence genes

A custom hybrid assembly script was used for hybrid *de novo* whole genome assembly, which differed from the pipeline used for SNP detection. ONT reads were basecalled and demultiplexed using Dorado V0.4.0 high accuracy model and reads with a Phred quality score of 8 or less were removed by chopper v0.5.0 (De Coster & Rademakers, 2023). Draft genome assemblies were generated from ONT reads using Flye v2.9.3 (https://github.com/mikolmogorov/Flye) (Kolmogorov et al., 2019) and polished with Medaka v1.8.1 (https://github.com/nanoporetech/medaka) using the same ONT reads. Illumina reads were paired and trimmed with BBDuk v39.0.1 (https://jgi.doe.gov/data-and-tools/software-tools/bbtools/bb-tools-user-guide/bbduk-guide/) and then mapped to the ONT draft assembly using BWA-MEM v0.7.18 (https://github.com/lh3/bwa). The resultant alignments were used to correct errors in the draft assemblies with polypolish v0.6.0 (https://github.com/rrwick/Polypolish). Mauve v2.4.1 (https://darlinglab.org/mauve/mauve.html) (Darling et al., 2011) aligned the *de novo* assemblies and genomes obtained from NCBI. Sequence reads and final assemblies were deposited into the Sequence Read Archive and NCBI GenBank, respectively (**Table 1**). Bakta 1.9.4 (https://bakta.computational.bio) was used for comprehensive taxon-independent annotation of coding sequences (CDS) (Schwengers et al., 2021). Bakta annotation results were screened based on a literature search of virulence factors in related *Mycoplasma* species using the UniProt protein product name identifier (Qin et al., 2019; Yiwen et al., 2021).

Candidate virulence genes from the annotated genomes were binned by function, and bins were grouped by virulence strategy, which included adhesion, invasion, evasion, biofilm formation, and cellular damage. The genes were then imported to a grouped table using GraphPad Prism v10.2.3, and a heatmap was generated from the dataset. The heatmap was annotated with Pixelmator Pro v3.6 (Pixelmator software, Vilnius, Lithuania). Biofilm formation factors were identified using BLASTp protein sequence similarity to well-characterized *M. gallisepticum* sequences (Ma et al., 2023; Wang et al., 2017; Xu et al., 2015).

## Results

### *In vitro* activity of antimicrobials against clinical isolates of *M. cynos* and *M. felis*

Super-modified Hayflick’s and SP4 broths were assessed for optimal growth of *M. cynos* and *M. felis* to prepare the inoculum of the CCU assay. Colour changes were observed for all *M. felis* and *M. cynos* isolates in SP4 broth, while in super-modified Hayflick’s broth, some of the isolates did not change the media colour (*M. cynos* n= 1*, M. felis* n= 3) (**Supplementary file 2**). Based on these results, SP4 broth was chosen to prepare the inoculum for the antimicrobial susceptibility assays.

All *M. cynos* (n=12) and *M. felis* (n=16) isolates were inhibited by the lowest tested concentration of tetracyclines, suggesting high *in vitro* susceptibility of *M. cynos* and *M. felis* to doxycycline, minocycline, and tetracycline (**Table 2**). The MIC values of fluoroquinolones varied from ≤ 0.125 µg/mL to 8 µg/mL. Still, the growth of most isolates was inhibited in the lowest tested concentrations of marbofloxacin, 0.125 µg/mL (*M. cynos* n = 6, and *M. felis* n = 6) and forenrofloxacin, 0.25 µg/mL (*M. cynos* n = 5, and *M. felis* n = 8). Isolates showed similar MICs to enrofloxacin and marbofloxacin, and this pattern was observed in both *M. cynos* and *M. felis* isolates (**Table 2**). The MICs of azithromycin (macrolide) for *M. felis* were observed in the highest concentrations tested (8 - 128 μg/mL), while MICs for most *M. cynos* isolates were observed within the lowest concentrations tested (0.25 - 1 μg/mL) (Table 2). The *M. bovis* P45 control strain presented a MIC range of 0.03 – 0.125 μg/mL for doxycycline, 4 μg/mL for azithromycin, 0.125 – 0.5 μg/mL for enrofloxacin, and ≤ 0.5 μg/mL for marbofloxacin. MIC values for all isolates are described in **Supplementary file 3**.

**Table 2.**
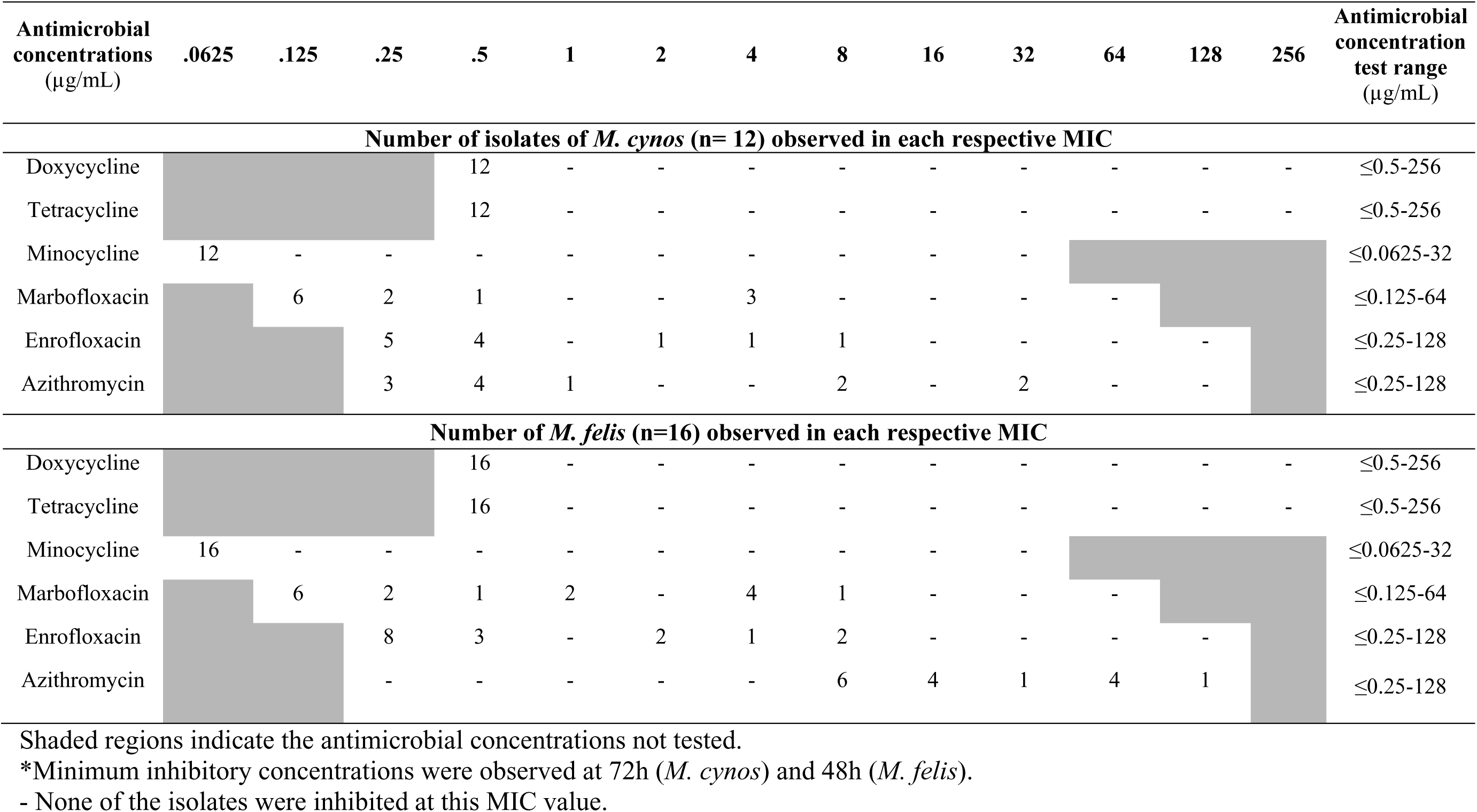
Distribution of the minimum inhibitory concentrations (MICs) obtained for *Mycoplasma cynos* and *Mycoplasma felis* clinical isolates by broth microdilution.

| Antimicrobial concentrations<br>(µg/mL) | .0625 | .125 | .25 | .5 | 1 | 2 | 4 | 8 | 16 | 32 | 64 | 128 | 256 | Antimicrobial concentration<br>test range<br>(µg/mL) |
| --- | --- | --- | --- | --- | --- | --- | --- | --- | --- | --- | --- | --- | --- | --- |
| Number of isolates of *M. cynos* (n= 12) observed in each respective MIC |  |  |  |  |  |  |  |  |  |  |  |  |  |  |
| Doxycycline |  |  |  | 12 | - | - | - | - | - | - | - | - | - | ≤0.5-256 |
| Tetracycline |  |  |  | 12 | - | - | - | - | - | - | - | - | - | ≤0.5-256 |
| Minocycline | 12 | - | - | - | - | - | - | - | - | - |  |  |  | ≤0.0625-32 |
| Marbofloxacin |  | 6 | 2 | 1 | - | - | 3 | - | - | - | - |  |  | ≤0.125-64 |
| Enrofloxacin |  |  | 5 | 4 | - | 1 | 1 | 1 | - | - | - | - |  | ≤0.25-128 |
| Azithromycin |  |  | 3 | 4 | 1 | - | - | 2 | - | 2 | - | - |  | ≤0.25-128 |
| Number of *M. felis* (n=16) observed in each respective MIC |  |  |  |  |  |  |  |  |  |  |  |  |  |  |
| Doxycycline |  |  |  | 16 | - | - | - | - | - | - | - | - | - | ≤0.5-256 |
| Tetracycline |  |  |  | 16 | - | - | - | - | - | - | - | - | - | ≤0.5-256 |
| Minocycline | 16 | - | - | - | - | - | - | - | - | - |  |  |  | ≤0.0625-32 |
| Marbofloxacin |  | 6 | 2 | 1 | 2 | - | 4 | 1 | - | - | - |  |  | ≤0.125-64 |
| Enrofloxacin |  |  | 8 | 3 | - | 2 | 1 | 2 | - | - | - | - |  | ≤0.25-128 |
| Azithromycin |  |  | - | - | - | - | - | 6 | 4 | 1 | 4 | 1 |  | ≤0.25-128 |

4 Shaded regions indicate the antimicrobial concentrations not tested.
\*Minimum inhibitory concentrations were observed at 72h (*M. cynos*) and 48h (*M. felis*).
6 - None of the isolates were inhibited at this MIC value.

### AMR-associated mutations in *M. cynos* and *M. felis* whole genomes

No SNP mutations associated with tetracycline resistance were observed in *M. cynos* and *M. felis* genomes corresponding to common resistance-conferring SNPs in *E. coli.* Further, there was also no evidence of *tet*(M) mediated resistance. No plasmids or other mobile genetic elements were found using the COPLA database.

For fluoroquinolones, mutations associated with resistance were found in *gyrA* for both *M. felis* and *M. cynos*; however, no mutations were detected in the *gyrB, parC,* or *parE* genes (**Table 3**). In *M. cynos,* 3/11 (27.3%) isolates had a mutation in Tyr154, including two Tyr154Gla and one Tyr154Ile. All three of these isolates showed elevated MICs to both marbofloxacin and enrofloxacin. The *gyrA* gene of *M. felis* was more polymorphic, with mutations detected at the Ser137, Glu141, and Asp150 loci. Two isolates (12.5%, 2/16) had Glu141Gly mutations, which were associated with elevated marbofloxacin and enrofloxacin MICs. Ser137Tyr and Asp150Asn mutations were present in three isolates (18.8%, 3/16), two of which contained both mutations and showed elevated marbofloxacin MICs.

**Table 3.** Single nucleotide polymorphisms (SNPs) associated with high minimum inhibitory concentration (MIC) of macrolide and fluoroquinolones in *Mycoplasma cynos* and *Mycoplasma felis* clinical isolates.

| Isolate ID | MIC (µg/mL) |  | Associated SNPs | MIC (µg/mL) | Associated SNPs |  |
| --- | --- | --- | --- | --- | --- | --- |
|  | Marbofloxacin | Enrofloxacin |  | Azithromycin | 23S rRNA (No. rRNA operons with mutation) | 50S L4 protein |
| ATCC | ≤ 0.125 | ≤ 0.25 | - | 0.5 | - | - |
| C01 | ≤ 0.125 | ≤ 0.25 | - | 0.5 | - | - |
| C02 | 0.25 | ≤ 0.25 | - | ≤ 0.25 | - | - |
| C03 | 4 | 8 | Tyr154Ile | 1 | - | - |
| C04 | ≤ 0.125 | 0.5 | - | 0.5 | - | - |
| C06 | ≤ 0.125 | ≤ 0.25 | - | ≤ 0.25 | - | - |
| C07 | 0.25 | 0.5 | - | 0.5 | - | - |
| C09 | 0.5 | 0.5 | - | 8 | - | - |
| C10 | ≤ 0.125 | 0.5 | - | 8 | - | - |
| C11 | 4 | 2 | Tyr154Ala | 32 | A2058G(2/3) | - |
| C12 | ≤ 0.125 | ≤ 0.25 | - | ≤ 0.25 | - | - |
| C14 | 4 | 4 | Tyr154Ala | 32 | A2058G(2/3) | - |
| F01 | ≤ 0.125 | ≤ 0.25 | - | 16 | - | - |
| F02 | 1 | 2 | - | 64 | C1187T(1/3) | Val43Leu |
| F03 | 4 | 8 | Ser137Tyr, Asp150Asn | 8 | C1187T(1/3) | - |
| F04 | ≤ 0.125 | ≤ 0.25 | - | 64 | - | Val43Leu |
| F06 | 4 | ≤ 0.25 | Ser137Ile, Asp150Asn | 8 | - | - |
| F07 | 8 | 4 | - | 16 | C1187T(1/3) | - |
| F08 | ≤ 0.125 | ≤ 0.25 | Ser137Ile | 8 | C1187T(1/3) | - |
| F09 | 4 | 2 | Glu141Gln | 16 | - | - |
| F10 | ≤ 0.125 | ≤ 0.25 | - | 8 | - | - |
| F11 | ≤ 0.125 | 0.5 | - | ≥ 128 | - | Val43Leu |
| F12 | 0.25 | 0.5 | - | 8 | - | - |
| F13 | 0.25 | 0.5 | - | 64 | - | Val43Leu |
| F14 | 0.5 | ≤ 0.25 | Asp150Asn | 8 | C1187T(1/3) | - |
| F15 | 4 | 8 | Glu141Gly | 32 | - | - |
| F16 | 1 | ≤ 0.25 | - | 64 | - | Val43Leu |
| F17 | ≤ 0.125 | ≤ 0.25 | - | 16 | C1187T(1/3) | - |

- indicates wildtype

For macrolides, we found multiple copies of the 23S rRNA gene in the investigated genomes. An A2058G substitution was identified in the 23S rRNA of *M. cynos* (2/12, 16.7%), which was correlated with azithromycin MICs of 32 μg/mL. A C1187T substitution was identified in 38% (6/16) of *M. felis* isolates, although we did not find any A2058G substitutions (**Table 3**). Elevated azithromycin MICs (≥8 µg/mL) were found for all *M. felis* isolates, and a novel 50S L4 ribosomal protein Val43Leu mutation was identified in *M. felis* (5/16, 31.3%) isolates, all of which also had azithromycin MICs ≥64 µg/mL.

### Putative virulence factors identified in *M. cynos* and *M. felis* whole genomes

A survey of peer-reviewed primary literature was conducted to determine virulence factors described in animal and human pathogenic *Mycoplasma* species (**Supplementary file 4**). Of the approximately 100 virulence factors described, 41 were identified in at least one *M. cynos* or *M. felis* whole genome through sequence homology or annotation name (**Figure 1**). Thirty uncharacterized lipoproteins were found by the Bakta annotation, which could not be further characterized by BLASTp or UniProt90 searches.

**Figure 1.**
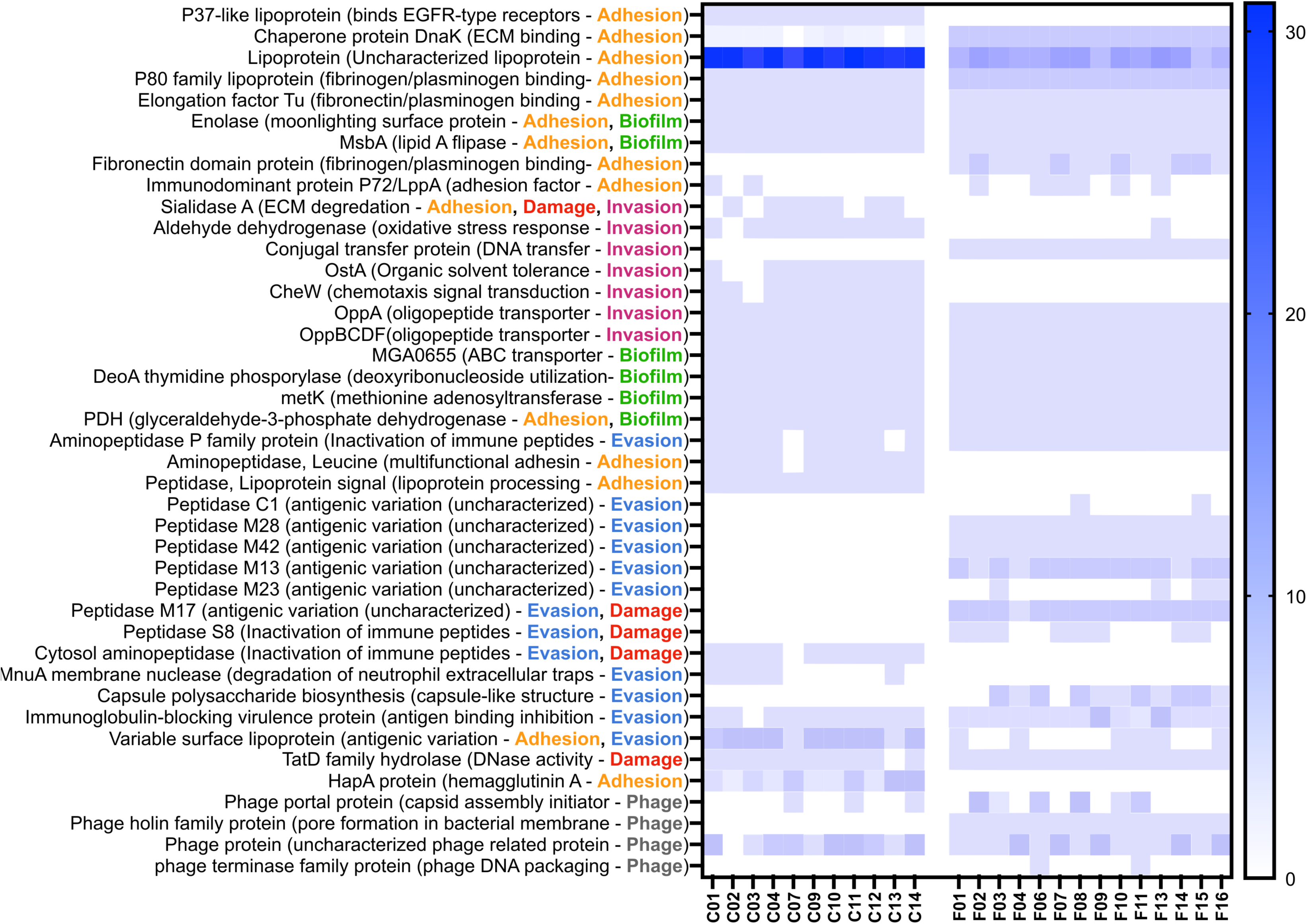
Heat map of virulence factors annotated by BAKTA from *Mycoplasma cynos* and *Mycoplasma felis* whole genomes isolated from dogs and cats with respiratory disease. **C**: *M. cynos*; **F**: *M. felis*.

Genes associated with adhesion and evasion covered most of the virulence factors identified, while genes for invasion and cellular damage were less numerous. *M. cynos* genomes contained more putative lipoprotein coding sequences than *M. felis*, however, these were numerous in both species. An assortment of peptidases responsible for post-translational modification of cell surface antigens were found primarily in *M. felis* isolates. Biofilm-related virulence factors were mainly associated with adhesion to host cell surfaces and were consistent across all the isolates (**Figure 1**). The following *M. gallisepticum* protein homologs were found in *M. cynos* and *M. felis* genomes: *msbA*, enolase, thymidine phosphorylase *DeoA*, methionine adenosyltransferase *metK*, glyceraldehyde-3-phosphate dehydrogenase and the *opp* operon, while protein-N(pi)-phosphohistidine-D-fructose phosphotransferase gene (*fruA*) was only identified in *M. felis* isolates (**Figure 1**). Other factors involved in *M. gallisepticum* biofilm formation could not be determined based on the predicted amino acid sequence in our isolates.

## Discussion

We investigated the *in vitro* activity of antimicrobials, resistance-associated mutations, and putative virulence factors in *M. cynos* and *M. felis* isolated from dogs and cats with respiratory disease, respectively. Key findings included that all isolates had low tetracycline MICs, and no 16S rRNA resistance determinants were identified, suggesting that tetracyclines remain an optimal antibiotic choice for treating *Mycoplasma*-associated respiratory infections in dogs and cats. Resistance-associated mutations were found in isolates with high MICs to fluoroquinolones and azithromycin. Further, most virulence factors identified were associated with adhesion and immune evasion.

We did not find evidence of tetracycline resistance mutations or *tet(M)*, and low MICs were observed in all isolates for tetracycline, doxycycline, and minocycline. Tetracycline resistance in *Mycoplasma* is commonly conferred by SNP mutations in the 16S rRNA gene in the tet-1 binding site formed by helix 34 between nucleotide positions 1054-1056 and 1196-1200 as well as helix 31 from 964-967 (Gautier-Bouchardon, 2018; Grossman, 2016; Sulyok et al., 2017). In *M. hominis*, ribosomal protection protein *tet(M)* was also associated with decreased susceptibility to tetracyclines; however, we did not find any evidence of plasmids in our isolates. Given the widespread use of doxycycline as a first-line antibiotic for canine and feline respiratory infections, it was unexpected to find consistently high susceptibility among our isolates (Lappin et al., 2017). These findings suggest that the sustained selective pressure from frequent tetracycline use does not drive resistance, and that tetracyclines remain a good choice for first-line treatment of *Mycoplasma* respiratory infections in dogs and cats.

Fluoroquinolone resistance is usually conferred by nucleotide substitutions in the quinolone resistance-determining region (QRDR) in the *gyrA*, *gyrB*, *parC*, and *parE* genes of DNA gyrase and topoisomerase IV enzymes, respectively (Bekő et al., 2020; Bokma et al., 2021; Sulyok et al., 2017). Ser83 and Glu87 mutations within the QRDR region have been reported in *M. gallisepticum*, *M. bovis*, *M. hominis*, and *M. genitalium* (Bokma et al., 2021; Chernova et al., 2016; Gautier-Bouchardon, 2018). The previously reported SNPs were mapped to Ser137 and Glu141 in our isolates. We identified *gyrA* mutations associated with increased fluoroquinolone MICs, including *M. cynos* Tyr154Ala/Ile, and *M. felis* Ser137Tyr/Ile, Glu141Gln/Gly, and Asp150Asn, which were within the *gyrA* QRDR. While these substitutions correlated with *M. cynos* MICs, they did not account for the phenotypic variability observed in *M. felis* isolates. We suspect SNPs in *parC* could also contribute to *M. felis* fluoroquinolone resistance, similar to findings from Bokma and colleagues (2021).

Macrolide resistance is associated with mutations in the 23S rRNA gene and amino acid substitutions in the L4/L22 ribosomal protein. An A2058G mutation in 23S rRNA, previously reported to be associated with macrolide resistance, was identified in *M. cynos* isolates that showed elevated MICs to azithromycin. Macrolide resistance in *Mycoplasma* spp. is associated with mutations in the peptidyl transferase loop of the 23S rRNA in domain V (*E. coli* positions 2000 to 2610) and domain II (*E. coli* positions 580 to 1250) (Bekő et al., 2020; Gautier-Bouchardon, 2018; Morozumi et al., 2005; Prats-van der Ham et al., 2018). Mutations at positions 2058, 2062, and 2611 are associated with macrolide resistance in different species of *Mycoplasma* (Kawai et al., 2013; Kong et al., 2016; Prats-van der Ham et al., 2018). The azithromycin susceptibility profile of *M. felis* was variable among isolates, and it was characterized by high MICs compared to *M. cynos*. The amino acid sequence of L4/L22 ribosomal protein is species-specific, making it challenging to compare SNP mutations across different *Mycoplasma* species (Khalil et al., 2017; Morozumi et al., 2005). We identified a novel mutation in the 50S L4 protein, specifically Val178Leu, in *M. felis* isolates with MICs greater than 64 µg/mL. Isolates that were only moderately susceptible had the C1187T mutation in the 23S rRNA and wild-type L4 protein. This suggests that the L4 mutation reduces azithromycin activity more than 23S mutations and that macrolide resistance results from multiple structures present at its binding site (Lovmar et al., 2009). Based on our *in vitro* and genomic findings, *M. cynos* and *M. felis* presented variable susceptibility to azithromycin, however, further studies are warranted to investigate its clinical efficacy.

Based on sequence homology to related *Mycoplasma* species, putative virulence factors associated with adhesion to host cells, immune system evasion, and biofilm formation were identified, suggesting that the virulence strategies of *M. cynos* and *M. felis* most resemble those of *M. hyopneumoniae* and *M. gallisepticum* (Jarocki et al., 2015; Leal Zimmer et al., 2020; Liu et al., 2019; Pflaum et al., 2018; Xu et al., 2015; Yiwen et al., 2021). Adherence factors are crucial for *Mycoplasma* attachment to host mucosal surfaces, with several homologs to the P1 adhesion complex of *M. pneumoniae* found in *M. cynos* and *M. felis* genomes. Moonlighting proteins like DnaK and elongation factor Tu also play critical roles in the adhesion process alongside P1 complex analogs (Dorigo-Zetsma et al., 2001; Widjaja et al., 2020). The major adherence strategies of *M. felis* and *M. cynos* likely use membrane-bound protein complexes that bind plasminogen and fibronectin of the host extracellular matrix. All *M. cynos* isolates contained one or more copies of *hapA*, a surface-presenting lipoprotein that binds canine erythrocytes (Kastelic et al., 2015). The oligopeptide permease (*oppABCDF*) operon, found in all *M. cynos* and *M. felis* genomes, governs the import of macromolecules and peptides (Wium et al., 2015). Our findings also indicate that immune evasion might be critical for the survival and pathogenicity of *M. cynos* and *M. felis*. The most prominent evasion strategy is likely through modulation of the host immune response using *Mycoplasma* immunoglobulin binding (MIB) proteins and *Mycoplasma* immunoglobulin proteases (MIP) (Arfi et al., 2016).

We based our search of biofilm-associated virulence factors on *M. gallisepticum* studies since its ability to form biofilms is well-documented in the literature (Chen, 2012; Ma et al., 2023; Wang et al., 2017). We identified multiple coding sequences previously demonstrated experimentally to be essential for biofilm formation in *M. gallisepticum*, including the *Opp* operon, which is required for transporting large molecules to the extracellular environment. We observed small cellular aggregates in liquid cultures of *M. cynos* and *M. felis*, demonstrating the precursor cell-cell attachment capabilities required for biofilm formation. These features could contribute to the invasion and persistence of these pathogens in host cell tissues, providing resistance to mechanical clearance by the innate immune system and increasing resistance to antimicrobials.

## Conclusion

This study lays the groundwork for establishing future clinical treatment guidelines, antimicrobial susceptibility testing, and enhanced diagnostic capabilities for *M. cynos and M. felis* respiratory infections. The demonstrated *in vitro* efficacy of tetracyclines supports their continued use as a first-line therapy in dogs and cats affected by *Mycoplasma*. Notably, we identified both established and potentially novel genetic markers linked to resistance mechanisms, including mutations in the 23S rRNA gene for macrolide and quinolone resistance-determining regions (QRDR) for fluoroquinolones. The identification of potential virulence genes associated with adhesion to host cells, immune system evasion, and biofilm provides a foundation for further research into *Mycoplasma* pathogenicity mechanisms. To our knowledge, this is the first study to provide in vitro susceptibility to antimicrobials and genomic analysis of AMR and virulence markers of *M. cynos* and *M. felis*.

## Data availability statement

The original contributions presented in the study are included in the article/supplementary material, further inquiries can be directed to the corresponding author. Hybrid assemblies obtained in this study can be found in the NCBI nucleotide database (Table 1).

## Author contribution

IF performed the experiment, conducted whole genome sequencing and assembly, and virulence analysis. MB supervised the project and assisted with the manuscript. PR performed the experiment, conducted SNP analysis, and revised the manuscript. SP and IM assisted with bioinformatic analysis and wrote the manuscript. JC supervised the project and assisted with the manuscript. GM conceived and designed the research, secured funding, supervised the project, and wrote the manuscript.

## Acknowledgements

The authors would like to thank Dr. Walter Demczuk for his assistance with the SNP analysis, and Dr. Hugh Cai for providing *M. cynos* and *M. felis* clinical isolates. We would also like to thank Jutta Hammermuller, Kai-Hsiang Chang and Karan Malhotra for their assistance with the MIC assays.

## List of Figures, Tables, and Supplementary Files

**Supplementary file 1.** Preparation of super-modified Hayflick’s broth.

**Supplementary file 2**. Color Changing Unit (CCU) assay results for *Mycoplasma cynos* and *Mycoplasma felis* clinical isolates using two different CCU methods.

**Supplementary file 3.** Minimum Inhibitory Concentration (MIC) values for clinical isolates of *Mycoplasma cynos* and *Mycoplasma felis*.

**Supplementary file 4.** Summary of the main virulence factors reported in pathogenic *Mycoplasma* species in animals and humans.

